# Anatomical structures, cell types, and biomarkers of the healthy human blood vasculature

**DOI:** 10.1101/2022.02.28.482302

**Authors:** Avinash Boppana, Sujin Lee, Rajeev Malhotra, Marc Halushka, Ellen M Quardokus, Bruce W. Herr, Katy Börner, Griffin M Weber

**Author notes:** Corresponding author Griffin M Weber.

## Abstract

More than 150 scientists from 17 consortia are collaborating on an international project to build a Human Reference Atlas, which maps all 37 trillion cells in the healthy adult human body. The initial release of this atlas provided hierarchical lists of the anatomical structures, cell types, and biomarkers in 11 organs. Here, we describe the methods we used as part of this initiative to build the first open, computer-readable, and comprehensive database of the adult human blood vasculature, called the Human Reference Atlas-Vasculature Common Coordinate Framework (HRA-VCCF). It includes 961 vessels and their branching connections, 10 cell types, and 10 biomarkers. With this paper we are releasing additional details on vessel types and subtypes, branching sequence, anastomoses, portal systems, microvasculature, functional tissue units, mappings to regions vessels supply or drain, and links to 3D reference objects. Future versions will add variants, geometric properties of vessels, and connections to the lymph vasculature; and, it will iteratively expand and improve the database as additional experimental data become available through the participating consortia.

## Background & Summary

We recently described an ongoing international effort from 17 consortia to construct a Human Reference Atlas (HRA) that maps the entire healthy adult human body down to the single-cell level^1^. It is a monumental task, considering the body has an estimated 37 trillion cells^2^. Combined, more than 150 experts worldwide are collaborating on this initiative. At the core of the HRA is a set of “ASCT+B” tables that contain hierarchical lists of *Anatomical Structures*, the *Cell Types* they contain, and associated *Biomarkers*. Many of the anatomical structures are linked to 3D reference objects. There are dozens of ASCT+B tables, each representing an organ or organ system. The completed HRA will encode the structure of tissues in the human body and their variability across individuals; it will increase the precision of physiological simulations; and, it will help researchers identify changes that occur during disease. The tables are created by an interdisciplinary team of domain experts who have, to-date, based the tables on existing knowledge, literature review, or experimental datasets. Over time, multimodal imaging and tissue assays applied to specimens being collected by the consortia will generate new knowledge that will be used to expand the ASCT+B tables and fill in details^3^.

One of the ASCT+B tables represents the blood vasculature. Blood vessels are both the source of life for people, bringing oxygen and nutrients to almost all living cells, as well as pathways that lead to disease, including coagulopathies in COVID-19, vascular abnormalities in diabetes, and the spread of metastatic cancers. For 2000 years, scientists have been cataloging different parts of the vasculature^4^, but to our knowledge, no one has yet connected the dots to create an open, computer-readable, comprehensive database of all the vessels throughout the healthy adult human body. This paper describes the methods we used to do this. The process involved creating a more extensive vasculature database, called the Human Reference Atlas-Vasculature Common Coordinate Framework (HRA-VCCF), with additional details beyond what is currently included in the ASCT+B tables. In this paper, we are releasing the first version of HRA-VCCF along with a database script used to derive the ASCT+B table from it.

Classic anatomy books, such as Gray’s Anatomy^5^ and Netter^6^, are not easily computer-readable, which prevents their use in software applications or linkage to biological datasets. The National Library of Medicine’s Visible Human Project created publicly-available anatomically detailed CT and MRI images^7^. However, these are not segmented into separate labeled structures. The Hemodynamics Modeling Laboratory group created a data repository of blood vessels and their geometric properties, such as length and diameter, for simulations^8^. However, this has not been released for public access. Terminologia Anatomica is an international standard for human anatomical structures, but its focus is on the naming of vessels, rather than their properties and relations to other structures, cell types, or biomarkers^9^.

The Uber anatomy ontology (UBERON) combines anatomical structures from different animal species, though an extract of just human structures exists^10^. Unfortunately, it is missing many blood vessels, especially related to microvasculature; many of the branching connections between vessels are also missing; and, because UBERON is a combination of different ontologies, there are inconsistencies in the level of detail provided across organs. The Foundational Model of Anatomy (FMA) ontology has a more complete list of blood vessels than UBERON, but it also has relatively little microvasculature^11^. There are again inconsistencies across organs, and the ontology is no longer being updated. Despite these limitations, the combination of UBERON and FMA was helpful to us in creating the HRA-VCCF blood vasculature database; and, we are working with UBERON to incorporate portions of the HRA-VCCF back into their ontology.

The blood vasculature is not only one of the organs included in the HRA, but it will also be used by the HRA as a coordinate system for describing the location within the body–a Vasculature Common Coordinate Framework (VCCF)^12^. Because the vasculature forms continuous pathways that extend to all living cells, it provides natural landmarks that scale to different body sizes and shapes. For example, a tissue specimen’s position can be recorded relative to nearby major arteries instead of an external reference point. In “genomic space”, a specimen can be described by the gene expression levels of blood vessel endothelial cells contained within that specimen, which show distinctive patterns depending on vessel type and part of the body^13,14,15^.

## Methods

### Blood vessel trees

The anatomical structures in the ASCT+B tables are meant to be a hierarchical “partonomy”. For other organs, anatomical structures are organized into a tree, with the “root” being the entire organ, the next level listing major structures contained within (i.e., “part_of”) the organ, the next level after that listing smaller subparts of those structures, and so on. The blood vasculature cannot be represented exactly this way because vessels are not contained within other vessels. As an analog, we used the chambers of the heart as the roots of four blood vessel trees: the left ventricle and right atrium for systemic arteries and veins, and the right ventricle and left atrium for pulmonary arteries and veins. The next level in the ASCT+B table lists the vessels that connect to the heart chambers, the level after that lists the vessels that branch off those vessels, and so on away from the heart and eventually extending out to the microvasculature.

As a starting point for creating these vascular trees, we selected 753 structures in UBERON that are subclasses of “blood vessel”. Many of these are actually groups of vessels, such as “arterial blood vessel” or “brain blood vessel”, rather than individual blood vessels. There are 29 relationship types between these structures in UBERON, with the most common being “is_a” relations–one vessel is within a larger grouping, such as “aorta” is a “thoracic cavity blood vessel”. For the ASCT+B tables, we were interested in physical connections between vessels, which use “branching_part_of”, “tributary_of”, and “part_of” relations. Of the 753 vessels, only 283 (38%) were connected to other vessels.

The data curation process began with AB and GW manually removing groups of vessels, as well as some non-human or developmental vessels. The remaining vessels were then connected to form the four trees. Because most vessels were not connected in UBERON, the missing connections had to be created. This often required also adding vessels from FMA that were missing from UBERON in order to complete the pathways back to the heart. Three domain experts then reviewed the initial trees: a cardiovascular pathologist (MH), a vascular biologist (RM), and a vascular surgeon (SL). They fixed mistakes and added vessels to expand details. Leveraging experts from three disciplines was helpful because they had complementary knowledge about different aspects of the vasculature. Of note, though, they had different preferred names for some vessels based on conventions in their respective disciplines. We generally used the clinical name for the vessel, though we might change this in the future.

A data cleanup step then corrected misspellings and ensured we named vessels in a consistent way. Then, duplicate vessels were merged. This occurred when a vessel is known by multiple names, and it was incorrectly represented in the trees as more than one vessel. Finally, we assigned a primary ID to each vessel, the ASID, which was based on the UBERON ID if possible. If the vessel was not in UBERON, we used an FMA ID if it existed.

While constructing the blood vessel trees, we also recorded several additional attributes about the vessels in the HRA-VCCF. First, we indicate the VesselType (chamber, artery, arteriole, capillary, venule, vein, sinus) and VesselSubType for capillaries (continuous, fenestrated, sinusoid). Then, we map each vessel to a BodyPart (e.g., kidney, liver, etc.) and BodySubPart. We flag vessels as belonging to a portal system or part of an organ’s functional tissue unit (FTU), note if they exist only in females or males, and indicate their BranchSequence (the order in which vessels branch from a parent vessel). Finally, we map major arteries and veins to anatomically correct 3D reference objects created by HuBMAP using data from the Visible Human female and male computed tomography datasets.

### Modeling approach

There were several challenges in representing the blood vasculature, which is really a network of vessels, as four trees. First, in UBERON there is a distinction between arteries branching into smaller vessels leading away from the heart and veins being treated as tributaries that merge into larger vessels leading back to the heart. This is not explicit in the ASCT+B table trees. In these tables, veins also start with the largest vessel and “branch” into smaller ones, opposite the direction of blood flow (Figure 1). However, the actual connection type is implicit based on the tree, with the left and right atrium trees corresponding to venous flow and tributary relations.

**Figure 1.**
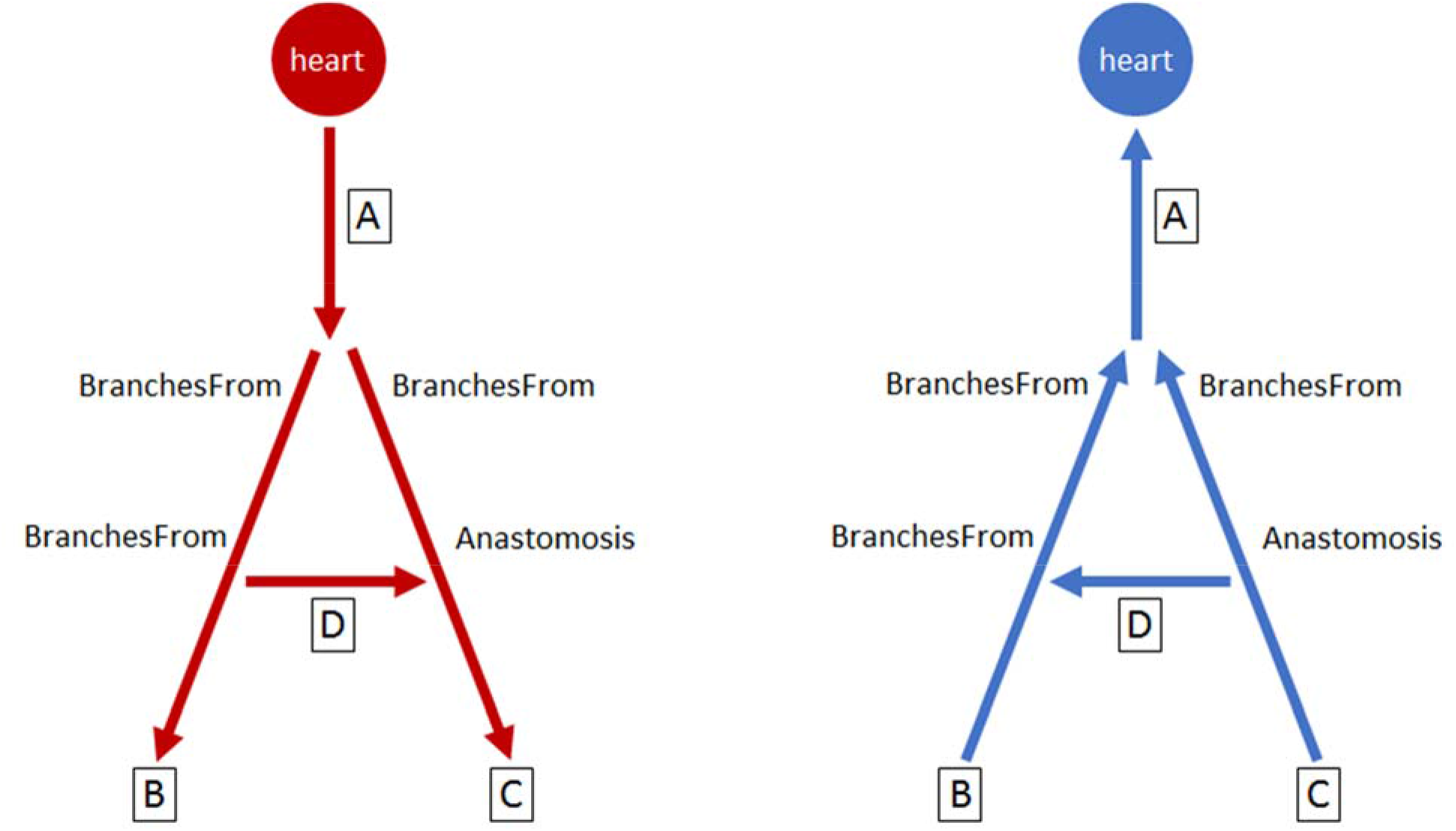
Blood vessel branching. Arrows point in the direction of blood flow. The end of a vessel that branches from another vessel (“BranchesFrom”) and anastomoses with another vessel (“Anastomoses”) is based on the direction of blood flow (arrows) and differs between arterial (left) and venous (right) vascular trees.

Next, anastomoses that form loops in the vasculature had to be broken. The branching side of the vessel was included in the ASCT+B table, while the anastomosis was recorded only in the HRA-VCCF. Sometimes the literature as well as our domain experts disagreed on which end of a vessel is the anastomosis. We decided to use the end where blood exits an artery or enters a vein as the anastomosis (Figure 1). An example is the anterior segmental medullary artery, which we listed as branching from the segmental spinal artery, rather than the anterior spinal artery, which we considered the anastomosis side. Blood direction in some vessels, such as sinuses in the brain, can change depending on the surrounding blood pressure. In these cases, we selected, to the best of our knowledge, the direction that is more common.

The sizes of the blood vessel trees can rapidly grow to a point where they become unmanageable if every vessel is explicitly included. To address this, we created “virtual vessels”, which represent a set of other vessels that have similar branches (Figure 2). For example, “renal artery” is a virtual vessel representing the left and right renal arteries. Instead of listing separate segmental renal arteries that branch off the left and right renal arteries, we only list them once as branches of the virtual renal artery. The virtual vessel itself branches off the same vessel as an arbitrarily selected vessel that it represents. For example, the renal artery branches off the abdominal aorta, just like the left renal artery.

**Figure 2.**
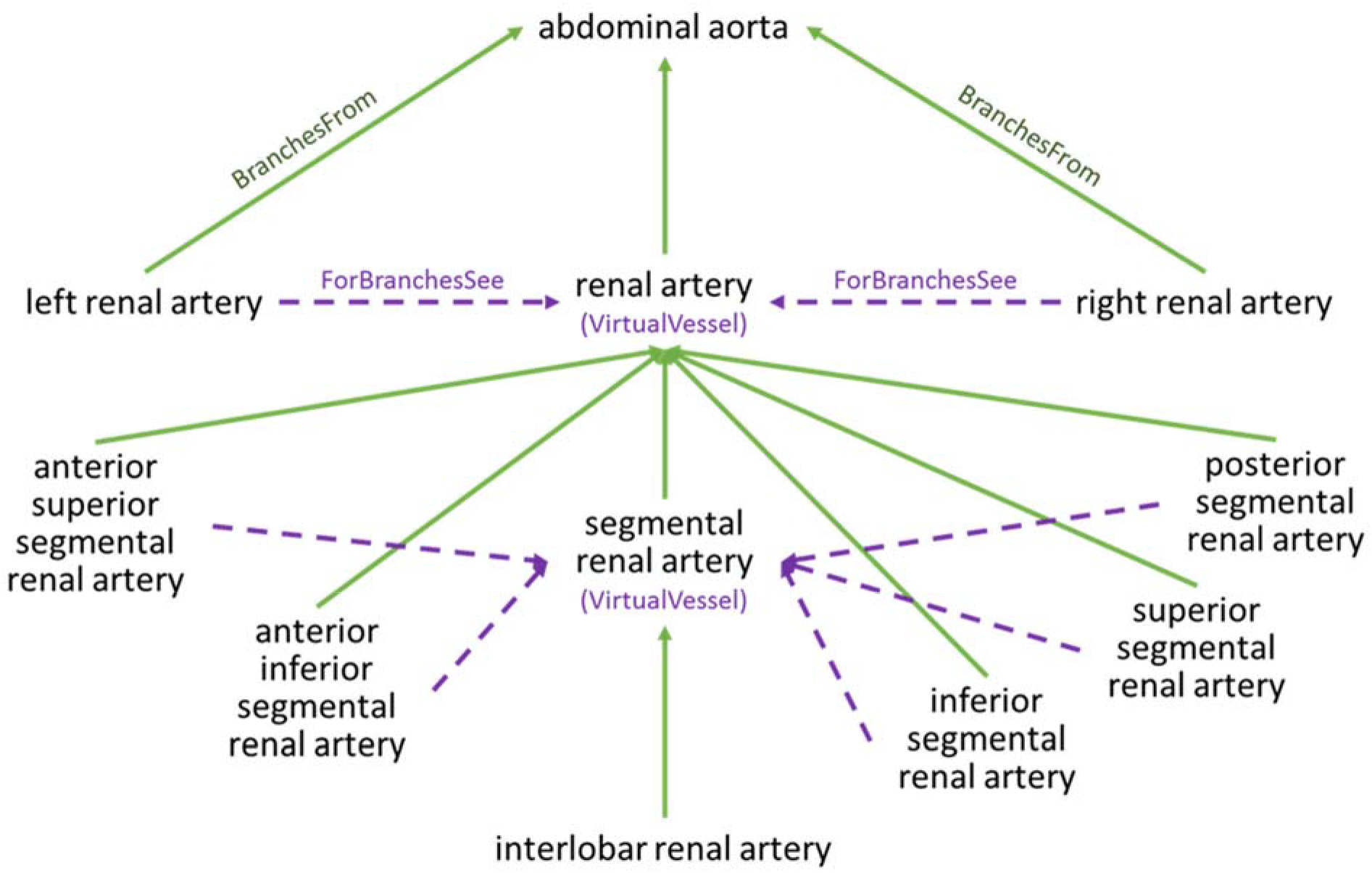
Virtual vessels. Virtual vessels simplify the vascular trees by enabling us to list the branches of a set of vessels only once. In this example of arteries in the kidney, interlobar renal artery is only listed once because both the left and right renal arteries and the multiple segmental arteries have been merged into virtual vessels.

This simplification of using virtual vessels treats sets of vessels as if they were identical, which is not truly the case. However, it might be sufficient for HRA applications. In future versions of our database, we can remove virtual vessels as needed if the branches of the different vessels need to be listed separately. Although it is not stated in the ASCT+B tables, the HRA-VCCF flags vessels as virtual vessels, and it points vessels being merged to their virtual vessel using a “ForBranchesSee” attribute.

Placing arteries and veins in separate trees for the ASCT+B table creates a problem in that it does not allow them to be connected through capillaries. We chose to include capillaries in the arterial trees (i.e., artery to arteriole to capillary) and end the venous trees at the venule level. In the HRA-VCCF, we use an “ArteryVeinConnects” attribute to link capillaries to their corresponding venules and venules to the corresponding capillaries. The database also includes an “ArteryVeinPair” for vessels such as the renal artery and renal vein, which do not directly connect, but have branches that do connect.

In portal systems, blood exits a capillary network and then enters another one before returning to the heart. We placed all the vessels of the hepatic portal system, including the veins draining into the hepatic portal vein and then leading into the liver, in the right atrium venous tree. However, in all other cases, we assigned both capillary networks and their connecting vessels to the left ventricle arterial tree (e.g., in the kidney, the glomerular capillary to the renal efferent arteriole to the peritubular capillary).

In some cases, there are multiple vessels with the same name that branch from different locations. These were distinguished in the ASCT+B table by adding “#N” to the vessel name. For example, the table lists both an “esophageal vein” branching from the azygos vein as well as an “esophageal vein #2” branching from the inferior thyroid vein. This is different from explicitly numbered vessels like “left posterior intercostal artery 3” and “left posterior intercostal artery 4”, which do not have a “#”. In the HRA-VCCF, the “VesselBaseName” attribute indicates the vessel name without the “#N” suffix.

### Cells and biomarkers

The ASCT+B tables list the cell types and biomarkers (CT+B) for each anatomical structure. Because of the large number of blood vessels, it was not practical to do this manually. Instead, we created a table in the HRA-VCCF with a set of “rules” that describe how to assign CT+B to vessels based on characteristics of those vessels. For example, one rule maps blood vessel endothelial cells to all vessels. Two additional rules map vascular associated smooth muscle cells and fibroblasts to all arteries and arterioles. Then we map as biomarkers (1) genes PECAM1 (CD31), CDH5 (CD144), TIE1, and TEK (TIE2) to blood vessel endothelial cells; (2) genes ACTA2, TAGLN (SM22), CNN1, MYOCD to vascular associated smooth muscle cells; and (3) AIFM2 (FSP1) and DDR2 to fibroblasts. We then created a database script that applies these rules sequentially, first to assign cell types to all vessels, and then to map biomarkers to vessels based on those cell types. Following HRA guidelines for the ASCT+B tables, each of the cell types used names and IDs from Cell Ontology^16^ and the biomarkers used names and IDs from the Human Genome Organisation (HUGO) Gene Nomenclature Committee (HGNC)^17^.

## Data Records

The HRA-VCCF contains 961 vessels, divided into four trees starting from the left ventricle, right atrium, right ventricle and left atrium, with 559, 361, 32, and 9 vessels, respectively (Table 1). The maximum depth of a vessel in the tree (distance from the heart) is 18, for the common plantar digital artery in the foot. It is reached through the following branching path: left ventricle > aorta > ascending aorta > aortic arch > descending aorta > descending thoracic aorta > abdominal aorta > common iliac artery > external iliac artery > common femoral artery > superficial femoral artery > popliteal artery > tibial-peroneal trunk > posterior tibial artery > lateral plantar artery > plantar arch > plantar metatarsal artery > common plantar digital artery. The ASCT+B table has an additional anatomical structure, “blood vasculature”, which serves as a parent node for the four trees. As a result, there are 18 + 1 = 19 levels in the ASCT+B table.

**Table 1.** Vessels by vascular tree

| Vascular Tree | Number of Vessels | Maximum Vessel Tree Depth | Mean Vessel Tree Depth | Vessels in UBERON (percent) | Vessels in FMA (percent) | Vessels in either UBERON or FMA (percent) |
| --- | --- | --- | --- | --- | --- | --- |
| left ventricle | 559 | 18 | 9.4 | 215 (38.5) | 467 (83.5) | 474 (84.8) |
| right atrium | 361 | 16 | 6.8 | 169 (46.8) | 309 (85.6) | 314 (87) |
| right ventricle | 32 | 7 | 4.8 | 4 (12.5) | 32 (100) | 32 (100) |
| left atrium | 9 | 5 | 2.6 | 2 (22.2) | 8 (88.9) | 8 (88.9) |
| Total | 961 | 18 | 8.3 | 390 (40.6) | 816 (84.9) | 828 (86.2) |

Table 2 shows the breakdown of vessels into seven vessel types and three sub-types. Table 3 and Table 4 list the ten functional tissue units^18^ and four portal systems modeled in this release. We mapped 71 vessels to female 3D reference objects and 76 vessels to male 3D reference objects (Figure 3). The database used 21 CT+B rules to assign 10 cell types and 10 biomarkers to vessels, for a total of 8,203 mappings.

**Figure 3.**
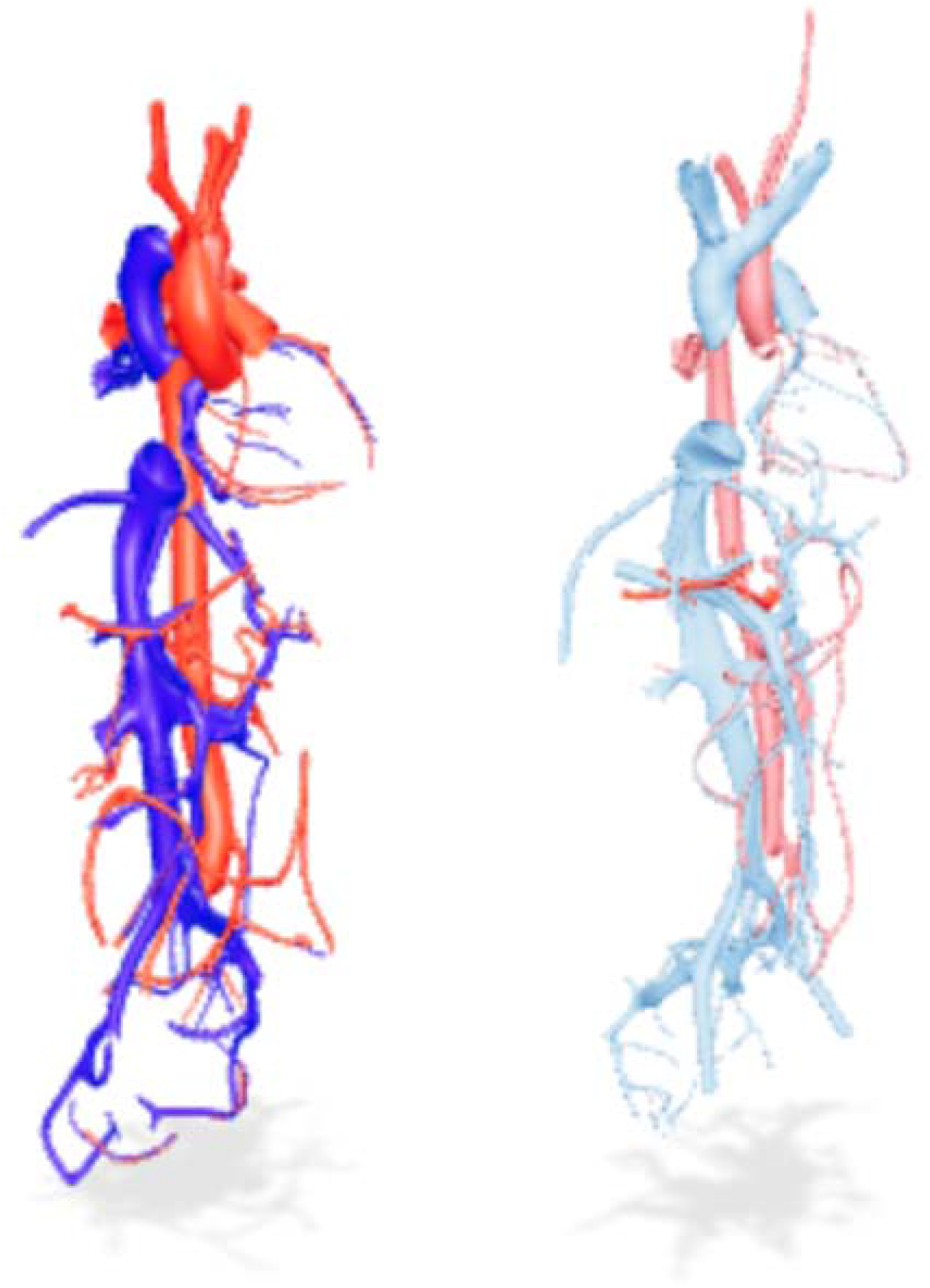
3D reference objects. We mapped major arteries and veins in the HRA-VCCF to anatomically correct 3D reference objects created by HuBMAP using data from the Visible Human female (left) and male (right) computed tomography (CT) datasets provided by the National Library of Medicine.

**Table 2.** Vessels by vessel type and subtype

| Vessel Type | Vessel Sub-Type | Number of Vessels |
| --- | --- | --- |
| heart chamber |  | 4 |
| artery |  | 563 |
| arteriole |  | 8 |
| capillary | continuous | 7 |
| capillary | fenestrated | 8 |
| capillary | sinusoid | 2 |
| venule |  | 5 |
| vein |  | 350 |
| sinus |  | 14 |

**Table 3.** Functional tissue units (FTU) included in the database

| Organ | Functional Tissue Unit | Number of Vessels |
| --- | --- | --- |
| kidney | long nephron | 1 |
| kidney | renal glomerulus | 1 |
| kidney | short nephron | 1 |
| large intestine | colonic crypt | 1 |
| liver | liver lobule | 3 |
| lung | alveolus of lung | 1 |
| pancreas | acinar cell of pancreas | 2 |
| pancreas | islet of Langerhans | 1 |
| spleen | red pulp of spleen | 1 |
| spleen | white pulp of spleen | 2 |

**Table 4.** Vascular portal systems included in the database

| Organ | Portal System | Number of Vessels |
| --- | --- | --- |
| liver and digestive system | hepatic portal system | 64 |
| kidney | renal portal system | 3 |
| pancreas | insulo-acinar portal system | 2 |
| pituitary gland | hypophyseal portal system | 4 |

**Table 5.** Selected columns in the Vessels table

| Field | Distinct values | Values, examples, or description |
| --- | --- | --- |
| VesselType | 7 | chamber, artery, arteriole, capillary, venule, vein, sinus |
| VesselSubType | 3 | capillaries: continuous, fenestrated, sinusoid |
| BodyPart | 49 | kidney, liver, brain, arm, leg, pelvis, abdominal cavity, etc. |
| BodySubPart | 10 | specific anatomic structure supplied or drained |
| PortalSystem | 4 | hepatic, hypophyseal, renal, insulo-acinar |
| Sex | 2 | female, male |
| Anastomoses | 37 | marginal artery of Drummond → branches of superior mesenteric artery |
| ArteryVeinConnects | 20 | liver arteriole ↔ hepatic sinusoid |
| ArteryVeinPairs | 365 | left renal artery ← . . . → left renal vein |
| ForBranchesSee | 152 | anterior superior segmental branch of right hepatic vein → segmental hepatic vein |
| VirtualVessel | 26 | segmental hepatic vein → represents 8 vessels |
| BranchSequence | 701 | the order in which vessels branch from a parent vessel |
| SpatialReferenceObjectFemale | 71 | pointers to 3D female reference objects |
| SpatialReferenceObjectMale | 76 | pointers to 3D male reference objects |

The HRA-VCCF is freely available as three data tables saved as comma-separated values (CSV) files with a Creative Commons Attribution 4.0 International (CC BY 4.0) license at https://github.com/GriffinWeber/HuBMAP/tree/main/HRA-VCCF

The Vessel table lists one row per blood vessel. The CellTypeBiomarker table lists “rules” describing cell types and biomarkers associated with different groups of vessels. The VesselCTB table is derived from the other two tables and lists the fully expanded mapping from vessel to cell types and biomarkers, once the rules are applied.

### Vessel table fields

Raw data fields:

- **BranchesFrom**. The “parent” vessel that is one step closer to the heart. For veins it is “drains to” rather than branches from.
- **Vessel**. The name of the blood vessel. The unique item (primary key) in this table. If a vessel has more than one BranchesFrom, the vessel is listed on multiple rows, but with “ #2”, “ #3”, etc. added to the end of its name.
- **VesselType**. Either heart chamber, artery, arteriole, capillary, venule, vein, or sinus.
- **VesselSubType**. For capillaries: continuous, fenestrated, sinusoid.
- **BodyPart**. A mapping from vessel to organ or part of the body.
- **BodySubPart**. The specific anatomical structure the vessel supplies or drains.
- **PortalSystem**. Indicates if the vessel is part of a portal system (e.g., hepatic portal system, hypophyseal portal system, etc.).
- **FTU**. Indicates if the vessel is part of a functional tissue unit (e.g., liver lobule, long nephron).
- **Sex**. Indicates whether the vessel is only found in males or females.
- **Anastomoses**. Indicates whether the vessel anastomoses with another vessel.
- **ArteryVeinConnects**. Indicates if an end branch (“leaf” vessel) in one vessel tree connects to a vessel in a different tree leading back to the heart (e.g., hepatic arteriole -> liver sinusoid).
- **ArteryVeinPair**. Indicates if another vessel has the same name, but with the words artery/arteriole swapped with vein/venule. Later this field will be used to match vessels with similar supplies/drains regions.
- **ForBranchesSee**. For some vessels, like the left and right renal arteries, rather than showing all the branches of both arteries, a “virtual” merged vessel is created (e.g., just “renal artery”). The branches are only added once to that virtual vessel. This field indicates the name of the virtual vessel that has the branches.
- **VirtualVessel**. This field contains a “1” if it is virtual merged vessel used to show the branches of other vessels; a negative value also indicates a virtual vessel, but the merged vessels are not yet explicitly defined in this table; zero for all other vessels.
- **BranchSequence**. The order in which vessels branch off of the BranchesFrom vessel. Vessels can have the same value if they branch off the BranchesFrom vessel at the same place. A value of 999 means the branching sequence will be added in a future version of this table.
- **SpatialReferenceObjectFemale**. The node name of the corresponding 3D reference object (female).
- **SpatialReferenceObjectMale**. The node name of the corresponding 3D reference object (male).
- **CoordX**. The X coordinate for the approximate location of selected vessels. For now, this is for debugging purposes, rather than to indicate the exact location.
- **CoordY**. The Y coordinate for the approximate location of selected vessels. For now, this is for debugging purposes, rather than the exact location.
- **ReferenceURL**. The website describing the vessel and where it branches from.
- **Reference**. A reference for the website. For example, if the ReferenceURL is a journal article, then this field is the reference (authors, title, journal, etc.) of the article.
- **ReferenceDOI**. The DOI of the reference if applicable.

Fields with values obtained from UBERON and FMA:

- **UBERON**. The ID of the vessel in the UBERON ontology.
- **FMA**. The ID of the vessel in the FMA ontology.
- **VesselTypeID**. The UBERON or FMA ID of the VesselType (imported from UBERON or FMA).
- **VesselSubTypeID**. The UBERON or FMA ID of the VesselSubType (imported from UBERON or FMA).
- **BodyPartID**. The UBERON or FMA ID of the BodyPart (imported from UBERON or FMA).
- **BodySubPartID**. The UBERON or FMA ID of the BodySubPart (imported from UBERON or FMA).
- **UBERONLabel**. The main label of the vessel in UBERON (imported from UBERON).
- **FMALabel**. The main label of the vessel in FMA (imported from FMA).
- **ASLabel**. The UBERON label of the vessel if it exists, otherwise the FMA label (imported from UBERON or FMA). AS = Anatomical Structure
- **ASID**. The UBERON ID of the vessel if it exists, otherwise the FMA ID (imported from UBERON or FMA). AS = Anatomical Structure
- **FTULabel**. The UBERON label of the FTU if it exists, otherwise the FMA label (imported from UBERON or FMA).
- **FTUID**. The UBERON ID of the FTU if it exists, otherwise the FMA ID (imported from UBERON or FMA).

Fields with values derived from the raw data:

- **VesselBaseName**. The vessel name without the “#N” at the end. This applies to vessels with more than one BranchesFrom.
- **VirtualVesselOfList**. The list (separated by semicolons) of vessels that are merged to form a virtual vessel.
- **VirtualVesselOfCount**. The number of vessels that are merged to form a virtual vessel.
- **VirtualInstances**. The number of times this vessel would be listed if virtual vessels were not used along the path back to the heart.
- **VirtualPath**. The list of vessels that have been merged along the path back to the heart.
- **PathFromHeart**. The list of branches leading from a heart chamber to the vessel. This field is useful for sorting the table.
- **PathFromHeartWithIDs**. Same as PathFromHeart, but the ASLabel and ASID are listed next to each vessel in the path.

### CellTypeBiomarker table fields

- **RuleID**. A unique ID for the row in the table, which also indicates the order in which the rules are applied.
- **MatchType**. The attribute used to select a group of vessels. For example, a list of vessel types (“VesselTypeList”). Used with MatchVal.
- **MatchVal**. The value of the attribute (from MatchType) used to select a group of vessels. For example, all arteries and arterioles (“artery;arteriole”).
- **IncludeCTB**. If “1”, then this rule defines a cell type (CT) or biomarker (B) that is associated with the matching vessels. If “0”, then this CT/B is NOT associated with the vessels.
- **CTBType**. If “CT1”, “CT2”, or “CT3”, then a hierarchical endothelial cell type of level N; if “CT" then a Level 1 non-endothelial cell type; if "BGene" then a gene biomarker.
- **CTBName**. The name of the cell type or biomarker.
- **CTBLabel**. The preferred label of the cell in Cell Ontology or the biomarker in HUGO Gene Nomenclature Committee (HGNC).
- **CTBID**. The ID of the cell in Cell Ontology or the biomarker in HUGO Gene Nomenclature Committee (HGNC).
- **ReferenceURL**. A website describing the cell type or biomarker.
- **Reference**. A reference for the website. For example, if the ReferenceURL is a journal article, then this field is the reference (authors, title, journal, etc.) of the article.
- **ReferenceDOI**. The DOI of the reference if applicable.

### VesselCTB table fields

- **Vessel**. The name of the blood vessel, linking back to the Vessel table.
- **BType**. The type of biomarker (e.g., BGene = gene biomarker).
- **CT1**. The name of the main cell type (level 1).
- **CT1Label**. The preferred label of the cell in Cell Ontology (level 1).
- **CT1ID**. The ID of the cell in Cell Ontology (level 1).
- **CT2**. The name of the cell sub-type (level 2).
- **CT2Label**. The preferred label of the cell in Cell Ontology (level 2).
- **CT2ID**. The ID of the cell in Cell Ontology (level 2).
- **CT3**. The name of the cell sub-sub-type (level 3).
- **CT3Label**. The preferred label of the cell in Cell Ontology (level 3).
- **CT3ID**. The ID of the cell in Cell Ontology (level 3).
- **BGene1**. The name of the gene.
- **BGene1Label**. The preferred label of the gene in HGNC.
- **BGene1ID**. The ID of the gene in HGNC.

## Technical Validation

### Source materials

The main references domain experts used to assemble and review the database are Gray's Anatomy^5^, Netter's Atlas of Human Anatomy^6^, Wikipedia^19^, and Radiopaedia^20^. Though numerous journal articles and other online resources were also used. These references are listed next to the corresponding vessel(s) in the database.

### Comparison to UBERON and FMA

Although we used UBERON as a starting point, only 390 (41%) of the final 961 vessels in our database are in UBERON. FMA was more complete, with 816 (85%) of our vessels in FMA; and, combined, 828 (86%) vessels had a match to either UBERON or FMA. Table 1 provides the breakdown by vessel tree. For the remaining 133 (14%) vessels without a match, the problem was often specificity. For example, FMA has one entity called “obtuse marginal artery”, while our database separates “obtuse marginal artery 1” and “obtuse marginal artery 2”.

In most cases we named vessels using the preferred label in UBERON or FMA. However, we used our own names in 233 (24%) vessels. (The UBERON and FMA IDs in our database enable users to link back to those ontologies to get the preferred label if needed.) Most of these were small modifications to provide consistency across all the vessels in the database. For example:

- We used singular forms of all vessel names, such as “long posterior ciliary artery” instead of “set of long posterior ciliary arteries”.
- We used consistent names for body parts, such as “interlobar renal artery” instead of “interlobar artery of kidney” to match other vessels containing the word “renal” instead of “kidney”.
- We specified the organ in vessels, such as “superior segmental artery of left lung” instead of “left superior segmental artery” to distinguish them from similarly named vessels in other organs.
- We replaced names of numbers with their equivalent numerals, such as “left posterior intercostal artery 3” instead of “left third posterior intercostal artery”. This was done because we envision the primary use case of this database to be computer-readable and used in other applications. Having numerals at the end of the name makes it easier for computer code to sort the vessels or remove the number.

### Limitations and future directions

This first release of the HRA-VCCF is meant to be the start, not the end, of an ongoing process that will iteratively improve and grow the database over time. The National Institutes of Health’s Human BioMolecular Atlas Project (HuBMAP) is funding this effort, which supports monthly meetings of an ASCT+B Working Group that reviews the status of each organ’s table, defines the format for future versions of the tables, prepares the tables for biannual production releases.

There are several limitations of this initial HRA-VCCF. It is not yet a complete list of all vessels in the human body. In many organs, based on the priorities of HuBMAP and other participating consortia, we went beyond what currently exists in UBERON. Though, much more work is still needed.

We included several examples of microvasculature in the database, especially where there are portal systems. However, the majority of arterial pathways are not yet connected to their corresponding veins. Microvasculature is more complex than major arteries and veins for several reasons. They are more often networks of vessels, which are harder to define as trees; there is very little literature about them, especially compared to larger vessels, probably because they are more difficult to study and less clinically important; descriptions of microvasculature are often simplified (e.g., “artery X ends in a capillary plexus”, leaving out the many small artery and arteriole branches); individual vessels at this level are not named, and terminology is inconsistent (e.g., “small vessels”, “arterioles”, “terminal branches”, “capillary network”, “capillary plexus”, “capillary bed”, etc.); and, it is often unclear from anatomical illustrations if thin wavy red and blue lines are real vessels or artistic interpretations of microvasculature. We plan to add new variables to the HRA-VCCF that capture characteristics of the microvasculature architecture, including the length and diameter of vessels, the branching depth, and branching angle. Prior work has shown that different tissues have distinctive microvasculature architecture patterns^21,22^.

The BodySubPart attribute of vessels will eventually map to specific structures in the other organ ASCT+B tables that the vessels either supply or drain. However, in this release we are only using BodySubPart to map a subset of microvasculature structures to functional tissue units. The BranchSequence orderings are not complete for all vessels.

The cell types and biomarker rules only cover a small number of well-known examples. As we expand this list in the future, for specific vascular cell types such as the endothelium and the vascular smooth muscle cell, some biomarkers are expressed ubiquitously in the vessels across all organ systems. However, further refinement is needed to identify biomarkers specific to vascular cells in individual organs; and, the use of large-scale single cell RNA sequencing, such as that offered by Tabula Sapiens^23^, is needed to advance our understanding of the organ-specific differences in vascular cells.

We did not include anatomical variants in this release. To the best of our knowledge, we used the most common arrangement of vessels. In the future, we plan to model variants similar to the CT+B rules, where each variant is a set of rules to remove structures in the reference vessel tree and add different ones. Where the data exists, we will indicate overall prevalence of a variant as well as breakdowns by subpopulations (e.g., sex and race). One challenge in performing the literature search to find this information is that there can be different naming conventions for the same variant^24^.

A separate parallel effort is constructing a similar ASCT+B table and extended database for the lymph vasculature. As that work progresses, we will link lymphatic vessels to corresponding blood vessels.

## Usage Notes

Our HRA-VCCF database is used to generate the blood vasculature ASCT+B table for the HRA. Releases of our HRA-VCCF database will occur at different times than its corresponding ASCT+B table, whose release dates are synchronized with the ASCT+B tables for other HRA organs. At the time of writing, the latest published blood vasculature ASCT+B table is version 1.1; while, the current HRA-VCCF database corresponds to blood vasculature ASCT+B table version 1.2, which is still in draft status.

The latest published ASCT+B tables, including the blood vasculature ASCT+B table, as well as standard operating procedures for constructing ASCT+B tables and an online tool to browse the tables (ASCT+B Reporter) are located on the HRA Portal, https://hubmapconsortium.github.io/ccf.

The draft blood vasculature ASCT+B table version 1.2 based on the HRA-VCCF data presented in this study is available at https://docs.google.com/spreadsheets/d/1pBO70FENOlSyPJukxHYjeMXW0SYTLj4lbcw2oGsjuf0/edit?usp=sharing.

Three-dimensional reference objects for selected blood vessels are available at https://hubmapconsortium.github.io/ccf/pages/ccf-3d-reference-library.html.

We welcome others to join our effort in constructing an HRA, and in particular helping with the blood vasculature ASCT+B table and HRA-VCCF. Signup for the ASCT+B Working Group is at https://iu.co1.qualtrics.com/jfe/form/SV_bpaBhIr8XfdiNRH.

## Code Availability

Three database scripts for Microsoft SQL Server are provided in GitHub at https://github.com/GriffinWeber/HuBMAP/tree/main/HRA-VCCF

The first file, HRA-VCCF_CreateTables.sql, creates three database tables to store the Vessel, CellTypeBiomarker, and VesselCTB data. The second file, HRA-VCCF_UpdateDerivedFields.sql, updates the derived fields in the Vessel table. This should be run if vessels are added or deleted or changes are made to the raw data columns. The third file, HRA-VCCF_GenerateExportTables.sql, applies the rules in the CellTypeBiomarker table to generate the VesselCTB data. It then outputs the data in ASCT+B table format and returns several aggregate summary tables.

## Acknowledgements

This research has been funded by the National Institutes of Health Human BioMolecular Atlas Program (HuBMAP) award OT2OD026671. We thank Zorina Galis for presenting us with the initial idea of using a Vasculature Common Coordinate Framework (VCCF) as a coordinate system for the human body and helpful suggestions and comments on constructing the HRA-VCCF database; Fauzan Isnaini for his contributions to modeling the vasculature in functional tissue units; and, Kristen Browne for creating the 3D reference objects.

## Author contributions

AB, MH, SJ, RM, GW performed data collection and curation; GW wrote the database software, performed data analysis, and wrote the original draft; all authors reviewed and edited the manuscript; and, KB obtained funding.

